# Information-based signal selection improves decoding of attention spotlight from multi-units & local field potentials and enhances correlation with behavior

**DOI:** 10.1101/2020.09.07.286195

**Authors:** C. De Sousa Ferreira, C. Gaillard, F. Di Bello, S. Ben Hadj Hassen, S. Ben Hamed

## Abstract

The ability to access brain information in real-time is crucial both for a better understanding of cognitive functions and for the development of therapeutic applications based on brain-machine interfaces. Great success has been achieved in the field of neural motor prosthesis. Progress is still needed in the real-time decoding of higher-order cognitive processes such as covert attention. Recently, we showed that we can track the location of the attentional spotlight using classification methods applied to prefrontal multi-unit activity (MUA) in the non-human primate (Astrand et al., 2016). Importantly, we demonstrated that the decoded (x,y) attentional spotlight parametrically correlates with the behavior of the monkeys thus validating our decoding of attention. We also demonstrate that this spotlight is extremely dynamic (Gaillard et al., 2020). Here, in order to get closer to non-invasive decoding applications, we extend our previous work to local field potential signals (LFP). Specifically, we achieve, for the first time, high decoding accuracy of the (x,y) location of the attentional spotlight from prefrontal LFP signals, to a degree comparable to that achieved from MUA signals, and we show that this LFP content is predictive of behavior. This LFP attention-related information is maximal in the gamma band. In addition, we introduce a novel two-step decoding procedure based on the labelling of maximally attention-informative trials during the decoding procedure. This procedure strongly improves the correlation between our real-time MUA and LFP based decoding and behavioral performance, thus further refining the functional relevance of this real-time decoding of the (x,y) locus of attention. This improvement is more marked for LFP signals than for MUA signals, suggesting that LFP signals may contain other sources of task-related variability than spatial attention information. Overall, this study demonstrates that the attentional spotlight can be accessed from LFP frequency content, in real-time, and can be used to drive high-information content cognitive brain machine interfaces for the development of new therapeutic strategies.

**Highlights:** We use machine learning to decode attention spotlight from prefrontal MUA & LFP.

We achieve high decoding accuracy of (x,y) spatial attention spotlight.

(x,y) attention spotlight position accuracy is maximal from LFP gamma frequency range.

MUA and LFP decoded attention position predicts behavioral performances.

Selecting high information signals improves decoding and behavioral correlates.

## Introduction

Accessing cognitive functions in real time, using machine learning methods applied to ongoing brain signals is considered as one of the major challenges of modern neurosciences, in order to enhance and restore human brain capacities (Astrand et al., 2014; Cinel et al., 2019; Dresler et al., 2018). Indeed, the ability to decode brain information in real-time is expected to allow for a better characterization of cognitive functions and development of therapeutic applications based on brain-machine interfaces. While great success has been achieved in the field of neural motor prosthesis (Prochazka, 2017), real-time decoding of higher-order cognitive processes such as spatial attention is still hampered by the complexity of these mechanisms.

One major issue in this respect is the fact that cognitive functions are mostly covert and can only be inferred transiently through subjects’ behaviors. Another crucial issue is the fact that cognitive processes are highly dynamic, irrespectively of behavioral goals or instructions (Gaillard et al., 2020).

In the last years, we have recorded multi-unit activity (MUA) signals from prefrontal frontal eye fields (FEF), a cortical region at the core of attention selection (Buschman and Miller, 2007; Ekstrom et al., 2008; Gregoriou et al., 2009; Ibos et al., 2013; Moore and Fallah, 2004; Wardak et al., 2006). We report real-time access to the (x,y) coordinates of attentional spotlight from these ongoing prefrontal neuronal population spiking activity, at high spatial and temporal resolution (Astrand et al., 2020; Di Bello et al., 2020; Gaillard et al., 2020). Importantly, we show a strong correlation between the decoded (x,y) attentional spotlight in real-time and subjects’ behavioral performance on a complex perceptual task.

In the following, we extend this (x,y) decoding of the attentional spotlight to local field potential (LFPs) signals, moving a step closer to real-time EEG based decoding of the attentional function. Indeed, LFP signals reflect the spiking activity that are summed over a large population of neurons while MUA refers to the activity of individual neurons or of a local population of neurons. While MUA activity is often best analyzed in the time-amplitude domain, LFPs are often analyzed in the time-frequency domain. Besides, we present a novel two-step decoding procedure optimizing correlation between decoded information and ongoing behavior. Specifically, we apply machine learning methods to neuronal population activities recorded from the FEF, bilaterally, while monkeys performed a cued spatial target detection task. We report for the first-time high (x,y) decoding accuracy of attentional spotlight location from LFP signals. We further show that LFPs attention-related informational content is maximal in the gamma frequency band. The real-time attention decoding accuracies for LFP are comparable to what we achieved from MUA and are highly correlated with behavioral performance. Based on the observation that the (x,y) attention spotlight location estimated from both MUA and LFP signals correlate with behavior, we introduce a novel attentional position decoding method based on a distinction between trials with high and low attention related information content. We demonstrate that this procedure improves decoding accuracies obtained from LFP and MUA signals and importantly, improves their correlation with behavior. This improvement is maximal for LFP signals compared to MUA signals, suggesting that LFP signals may contain other sources of task-related variability than spatial attention information, a point that is highly relevant for attentional processes decoding. Overall, this study provides methodological bases to drive high attention-information content cognitive brain machine interfaces from both MUA or LFP activities. It also opens the way to targeting other cognitive functions such as working memory, and possibly extend this approach to non-invasive signals such as EEG or fMRI signals.

## Methods

### Subjects and surgical procedures

Two adult male rhesus monkeys (Macaca mulatta) were used in this experiment. All surgical and experimental procedures were approved by the local animal care committee (C2EA42-13-02-0401-01) in compliance with the European Community Council, Directive 2010/63/UE on Animal Care. The surgical procedures, the FEF location, and visual stimulation techniques have been described elsewhere (Astrand et al., 2016).

### Behavioral task

The task is a 100% validity endogenous cued spatial target detection task (fig. 1A). The animals were placed in front of a PC monitor (1920×1200 pixels and a refresh rate of 60 HZ), at a distance of 57 cm, with their heads fixed. The stimuli presentation and behavioral responses were controlled using Presentation (Neurobehavioral systems®, https://www.neurobs.com/). To start a trial, the bar placed in front of the animal’s chair had to be held by the monkeys, thus interrupting an infrared beam. The onset of a central blue fixation cross (size 0.7°×0.7°) instructed the monkeys to maintain eye position inside a 2°×2° window, defined around the fixation cross. To avoid the abort of the ongoing trial, fixation had to be maintained throughout trial duration. Eye fixation was controlled thanks to a video eye tracker (Iscan(tm)). Four gray square (size 0.5°×0.5°) were displayed, all throughout the trial, at the four corners of a 20°x20° hypothetical square centered onto the fixation cross. Thus, the four squares (up-right, up-left, down-left, down-right) were placed at the same distance from the center of the screen having an eccentricity of 14° (absolute x-and y-deviation from the center of the screen of 10°). After a variable delay from fixation onset, ranging between 700 – 1200 ms, a small green square, the cue (size 0.2°×0.2°) was presented, for 350 ms, close to the fixation cross (at 0.3°) in the direction of one of the grey squares. Monkeys were rewarded for detecting a subtle change in luminosity of this cued square - i.e., the target. The change in target luminosity occurred unpredictably between 350 to 3300 ms from the cue off time. In order to receive a reward (drop of juice), the monkeys were required to release the bar in a limited time window (150 - 750 ms) after the target onset (hit trials). In order to make sure that the monkeys did use the cue instruction, on half of the trials, distractors were presented during the cue to target interval. Two types of distractors could be presented: (i) an uncued distractor (33% of trials with distractor) - that could take place equiprobably at any of the uncued locations; (ii) a workspace distractor (67% of trials with distractor) - that correspond to a small square presented randomly in the workspace defined by the four target locations. The contrast of the square with respect to the background was the same as the contrast of the target against the grey square. The monkeys had to ignore all of these distractors. Responding to any of them interrupted the trial. If the response occurred in the same response widow as for correct detection trials (150 - 750 ms), the trial was counted as a false alarm (FA) trial. Failing to respond to the target (Miss) similarly aborted the ongoing trial. Overall, data was collected for 19 sessions (M1: 10 sessions, M2: 9 sessions). The behavioral performance of each animal is presented in figure 1B (proportion of hits over miss and FA trials).

**Figure 1:**
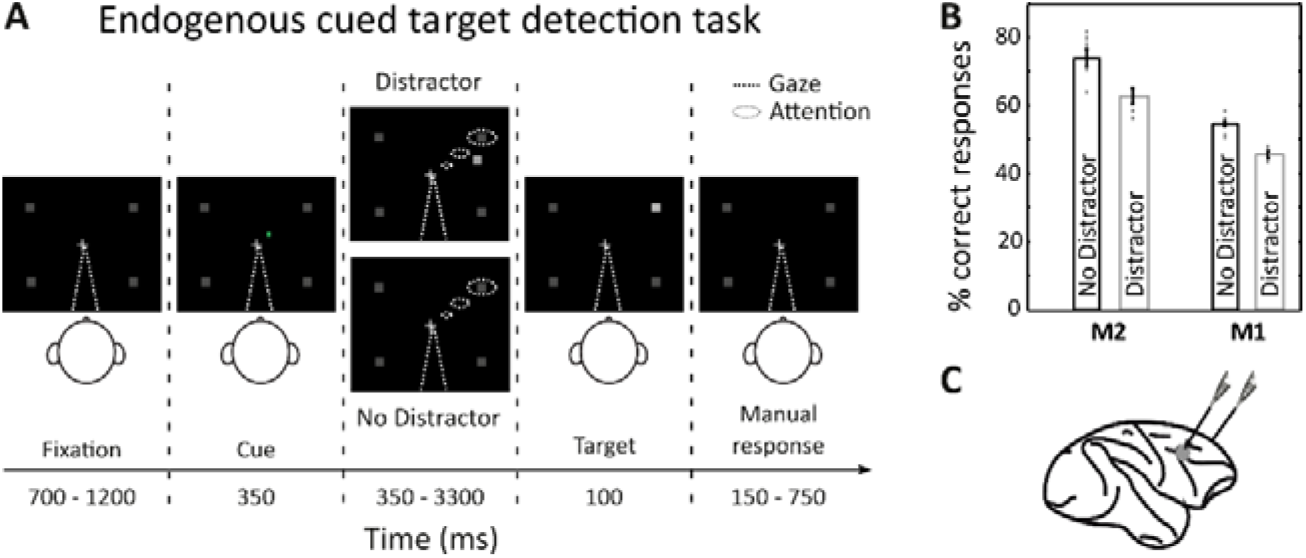
Task design and behavioral performance. (A) 100% validity cued spatial target detection task with distractors. Monkeys had to hold a bar and fixate a central cross on the screen for a trial to be initiated. Monkeys received a liquid reward for releasing the bar 150 - 750 ms after target presentation. Target location was indicated by a cue (green square, second screen). Monkeys had to ignore any uncued event. (B) Behavioral performance of monkeys M1 and M2 at detecting the target in the presence or absence of a distractor (median % hits +/-median absolute deviations, dot correspond to individual sessions). (C) Recording sites. On each session, 24-contact recording probes were placed in each FEF.

### Recording techniques

Bilateral simultaneous recordings in the two frontal eye fields (FEF) were carried out using two 24-contact Plexon U-probes (fig. 1C). The contacts had an interspacing distance of 250 μm. Neural data was acquired with the Plexon Omniplex® neuronal data acquisition system. The data was amplified 400 times and digitized at 40.000 Hz. A threshold defining the multi-unit activity (MUA) was applied independently for each recording contact before the actual task-related recordings started. Local field potentials (LFP) were recorded simultaneously on the same electrodes as MUA. The neuronal properties of the recorded neuronal sample have already been described elsewhere (Astrand et al., 2020; Gaillard et al., 2020).

### Neuronal decoding procedure

MUA and LFP signals were aligned on the target presentation time and sorted according to the monkey’s behavioral response (hits and misses). Fast Fourier transform analyses were performed on LFP signals recorded on all 48 channels to quantify signal power up to 250 Hz. Signal normalization was applied to LFP signals.

Specifically, instantaneous powers were z-scored with respect to a pre-cue baseline, by subtracting from these instantaneous power frequency series the power average of the 300 ms before the cue presentation and dividing by the standard deviation of this signal over all trials. Decoding from LFP signals was then performed either on unfiltered data or on eight independent frequency bands: δ (0-4 Hz), θ (4-8 Hz), α (8-12 Hz), low β (12-20 Hz), high β (20-30 Hz), low γ (30-60 Hz), mid γ (60-120 Hz), high γ (120-250Hz). As in Astrand et al., (2016, 2015), a regularized linear decoder was used to associate, on hit trials, the neuronal activity associated to one of the four possible target locations.

Decoder input signals corresponded either to the number of spikes for MUA or to the normalized instantaneous power of all frequencies or specific frequency bands for LFPs, computed over the specified time window. On each given time interval before target presentation, the decoder was trained on a random set of 70% of hit trials and then tested on the 30% remaining hit trials and all misses, with activities sampled at the same interval as the training interval. Trial positions were equalized in the training set to avoid any decoding bias. To avoid overfitting, training and testing were performed from different trials. During training, the input of the classifier was a 48-channel by N-trial matrix, corresponding to the average neuronal signal computed over the time interval of interest, for each of the 48 recording channels and each of the N training trials. As an input, the decoder also used the (x,y) coordinates of the target for each of these N training trials. During testing, for each trial, new to the classifier, the output of the classifier was estimated from a 48-channel vector corresponding to the average neuronal signal on the time interval of interest, on each of the 48 recording channels, on the considered testing trial. The output calculated by the decoder correspond to an X and Y coordinate. Thus, it could be read as an (x,y) estimation of attentional spotlight or as a quadrant category, corresponding to one of the four possible target localization (as in Astrand et al., 2016, 2015, 2014; Gaillard et al., 2020). Training and testing were performed on neuronal signals from 10 ms to 1200 ms before target presentation with a time step of 20 ms. All trials with cue-to-target intervals shorter than 1700ms were excluded from this analysis. For each interval, training and testing steps were repeated 100 times, then averaged to define a decoding performance corresponding to the number of correct classifications according to quadrant categories. We estimated the 95% confidence interval to verify the statistical significance of our decoding performance. The same decoding analyses as described above were used with a training set based on random labels. In other words, the decoder used the same neuronal signal, but the coordinates of the target were randomized and thus did not correspond to the actual condition in which the neuronal signal was recorded.

### Behavioral correlation

In order to validate the decoding procedure, we investigated the correlation between the (x,y) attentional spotlight decoded from neuronal signals with monkey’s behavioral response (Percentage of hits over miss trials). Specifically, the relative distance between the actual target location and the decoded attentional spotlight location was calculated for each trial. Percentage of hits over miss trials was then calculated over 0.5° distance vectors. To avoid biases, total number of hits and misses were equalized and then binned - the whole procedure was repeated 100 times. The X and Y location of attentional spotlight was calculated from a leave-one-out decoding strategy (i.e., training was performed on all hit trials except one used for the testing). For misses, the decoder was trained on all hit trials and tested on all misses. Training and testing were performed on a 150 ms time window prior to target presentation. Statistical analyses were carried using linear regression model.

### Two-step decoding procedure

In this part, we dissociate high attention-related informational spatial content trials from low attention-related informational content trials. We use the relative distance calculated between the decoded (x,y) attentional spotlight (AS) and the real target location (T) for hit trials, as described above. Two categories of hit trials were identified from this first decoding: 1) trials in which the decoded attentional spotlight is close to target location (i.e. HighContent trials) and 2) trials in which the decoded attentional spotlight is far from target location (i.e. LowContent trials). HighContent and LowContent trials were defined according to a threshold of 7° between real target location and decoded attentional spotlight (HighContent trials: |AS-T|<7°; LowContent trials: |AS-T|>=7°). Given the high difficulty of the task, monkeys cannot succeed in the trial if they are not orienting their attention near to the target location (Astrand et al., 2016). Thus, we hypothesized that these differences between HighContent and LowContent trials was due to differences in spatial attention informational content between these two types of hit trials, and that signals were more rpresentative of the expected target location in HighContents trials that in LowContent trials. Decoding performance and behavioral correlation were thus calculated a second time as follows. In order to evaluate classification performance, training was performed on all HighContent trials and testing was performed on different percentages of HighContent trials over LowContent trials (0% to 100% ratio). The proportion 70/30 of trials used for training and testing was conserved. Once training and testing sets were selected, the decoding procedure applied was the same that the procedure described in the previous section. In order to evaluate the correlation between decoded attention position and behavioral performance, we performed a trial by trial (x,y) estimation of attentional position. More specifically, for HighContent trials position decoding, the decoder was trained on all HighContent trials except one and tested on the remaining one (leave one out strategy). For LowContent trials and misses, the decoder was train on all HighContent trials and tested on LowContent trials and misses. Training and testing were performed 150 ms before target presentation. The relative distance between AS and T was calculated and associated with the percentage of hit trials with respect to misses. Hit trials included 50% of LowContent trials and 50% of HighContent trials. For each signal (MUA and LFP), we compared the effect of HighContent trials on decoding performance and behavioral correlation. Statistical comparisons were performed using non parametric tests (Wilcoxon rank sum test) and multiple linear regressions.

## Results

In order to access the location of the attentional spotlight, a linear decoder was used to estimate the (x,y) coordinates of attention based on MUA and LFP signals, recorded from the prefrontal cortex (FEF, bilaterally, Fig. 1C) while monkeys performed a cued target detection task (Fig. 1A). The readout of this linear decoding procedure can be classified in one of four possible classes indicating whether attention is correctly oriented to the cued visual quadrant (correct classification), or to one of the three other quadrants (incorrect classification, Astrand et al., 2014, Tremblay et al., 2015b). Alternatively, the readout of the linear decoding can be taken as an error to the cued location and transformed into an (x,y) continuous coordinate (Astrand et al., 2020, 2016; Gaillard et al., 2020). In the first part of the results, we report for the first time continuous attentional spotlight position decoding from LFP signals, with performance accuracy levels similar to MUA based decoding. We then analyze how the continuous (x,y) estimates of attentional spotlight based on prefrontal MUA and LFP signals predict behavioral performance, thus validating the decoding procedure. Finally, we develop a decoding method that optimizes the spatial decoding of attention from MUA and LFP signals and highlights qualitative variability in prefrontal attention related information.

### Classifying spatial attention from prefrontal MUA and LFP

Figure 2A and 2B represent the classification performance based respectively on FEF recorded MUAs and LFPs (irrespective of frequency content). Neuronal activity (decoder input) was averaged just prior to target presentation, calculated across varying time windows ranging from 10ms to 1200ms. Decoding accuracy on hit trials is significantly higher than chance for both MUA (Fig. 2A, blue, mean = 77%, s.e. = 2.1%, for window size = 1200ms, dashed blue line, 95% C.I, note that absolute chance level is at 25%) and LFP signals (Fig. 2B, blue, mean = 71%, s.e. = 2.1%, for window size = 1200ms, dashed blue line, 95% C.I). Thus, on hit trials, spatial attention can be successfully classified from both MUA and LFP signals.

**Figure 2:**
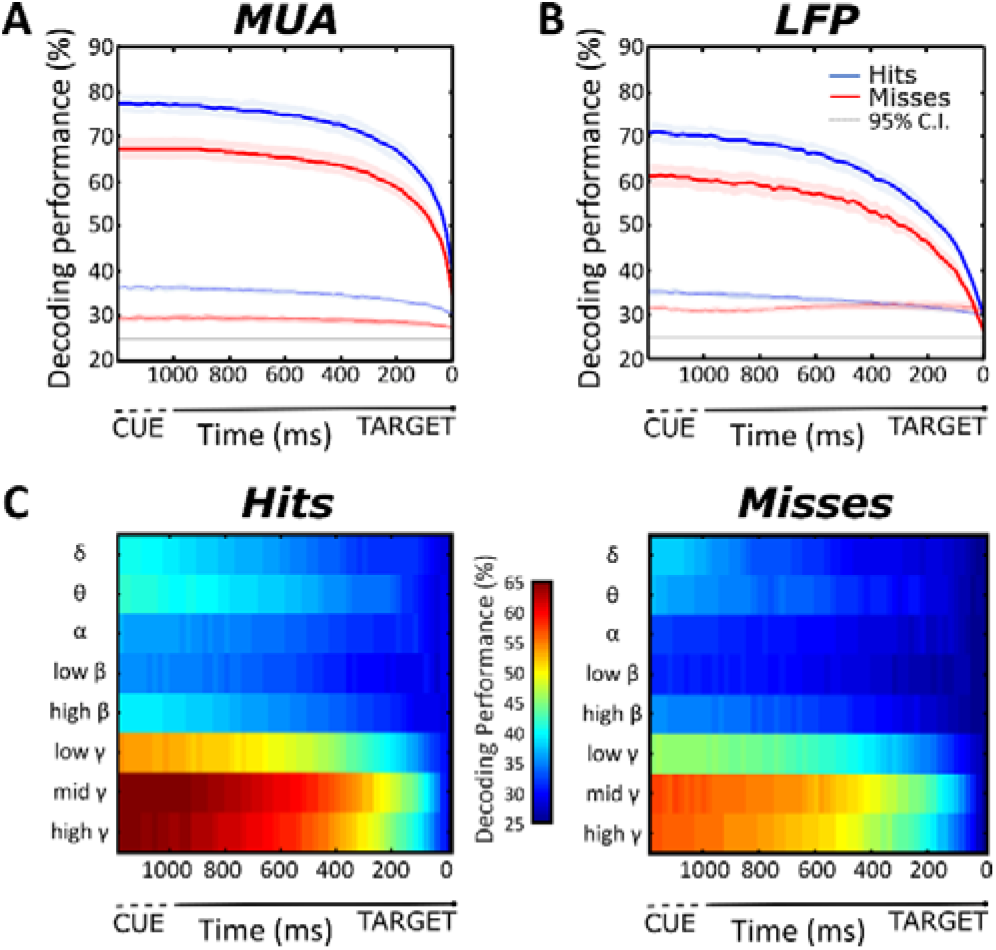
Spatial attention decoding accuracies from (A) multi-unit activity (MUA) or (B) local field potentials (LFP), as a function of averaging time window size from target onset (0 ms), on hits (blue, mean +/-s.e.) and miss trials (red, mean +/-s.e.). Black dashed line (25%): absolute chance level; dashed blue and red curves: 95% C.I. for hit and miss trials. (C) Spatial attention decoding accuracy from LFP signals per LFP frequency band, as a function of averaging time window size from target onset (0 ms), on hit (left) and miss trials (right).

Interestingly, decreasing time intervals before target presentation highly impacts decoding accuracies. Performances decrease from 77% to 40% for MUA (Fig. 2A, blue, mean =40%, s.e. = 1.7%) and from 71% to 29% for LFPs (Fig. 2B, blue, mean = 29%, s.e. = 0.9%). Additionally, for LFP signals, classification significance is reached only for window sizes starting from 30 ms (Fig. 2B, blue, mean=29%, s.e.=0.9%). Compared to short time windows, longer time windows reflect average spatial attention location, and thus yield higher classification rates. On both signals, window size thus implies a trade-off between temporal resolution and overall classification accuracy.

While MUA signals are processed in the time-amplitude domain, LFP signals are processed in the time-frequency domain. In the following part, we segregated the different functional frequency bands of LFPs to investigate their specific impact on classification performances. Figure 2C represents the decoding accuracy in time as a function of specific LFP functional frequency band content. As observed on the overall decoding accuracy from all LFP frequency content decomposition, larger window sizes yield higher decoding accuracies in all frequency bands (Fig. 2C). However, information about spatial location of attention is mainly contained in the gamma frequency bands (30-250 Hz). Specifically, on hit trials, for the largest window sizes, decoding accuracies are below 50% for all frequency bands <30Hz (δ = 40%; θ = 42%; α = 36%; low β= 35%; high β= 39%) and reach a maximum of 54% for low γ (30-60 Hz), 66% for mid γ (60-120 Hz) and 65% for high γ (120-250Hz) (Fig.2C). In addition, full spectrum LFP decoding accuracy is higher compared to LFP band-specific decoding accuracies.

For both MUA and LFP signals, decoding is significantly more reliable on hit trials than on misses at all window sizes (e.g. window size = 1200ms, MUA: Fig. 2A, blue, mean = 77%, s.e. = 2.1% vs. red, mean = 67%, s.e. =2.4%; LFP: Fig. 2B, blue, mean = 71%, s.e. = 2.1% vs. red, mean = 61%, s.e. = 2.4%). This holds true for all LFP frequency bands, although impact of negative trial outcome is stronger on higher LFP frequency bands as compared to lower (Fig. 2C). Overall, this supports that spatial attention is miss allocated during miss trials (Astrand et al., 2016, Gaillard et al., 2020), subsequently interfering with perception (Astrand et al., 2020).

### The decoded (x,y) attentional spotlight predicts behavior

Spatial attention is a covert cognitive process. Therefore, it is not possible to verify spatial attention decoding behavioral significance by a direct single trial correlation between decoding readout of spatial attention and an observable behavioral measure other than target detection performance. It is however possible to validate the correlation between the decoding readout of spatial attention and behavior over multiple trials (Astrand et al., 2016, Gaillard et al., 2020). In the following, attention to target distance is defined, in each trial, as the distance between expected target location and the corresponding decoded (x,y) attentional spotlight, 150 ms before target onset. This distance parameter is then correlated to a behavioral performance calculated as the percentage of hits over miss trials. For all signal types, we observe that monkeys produce more hits when their attentional spotlight is deployed closer to target location. Specifically, we demonstrate a significant linear correlation between the distance of decoded attentional spotlight to target and the hit rate, when using MUA based decoding (Fig. 3A. linear regression: r^2^ = 0.48, F= 86, p-value < 0.005) as well as when using LFP based decoding (Fig. 3.B linear regression: r^2^= 0.65, F= 174, p-value < 0.005). This indicates that similarly to MUAs, LFPs spatial attention information predicts behavior.

**Figure 3:**
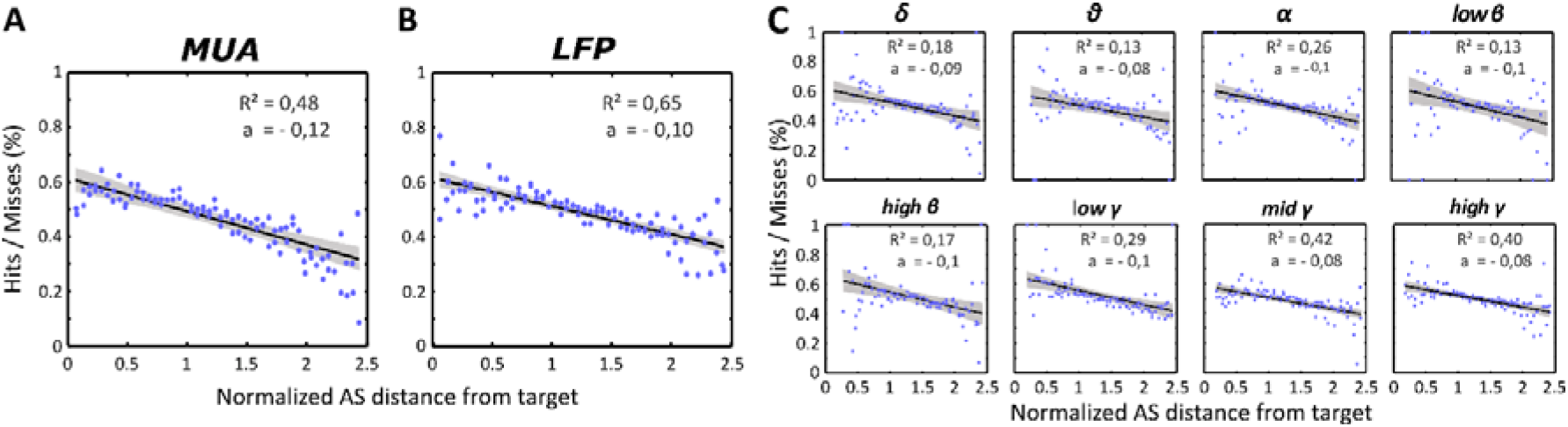
Correlation between behavioral performances & distance between the attentional spotlight and the target location from (A) multi-unit activity (MUA), (B) local field potentials (LFP), on all frequency power content or (C) as a function of specific frequency ranges ((δ (0-4 Hz), θ (4-8 Hz), α (8-12 Hz), low β (12-20 Hz), high β (20-30 Hz), low γ (30-60 Hz), mid γ (60-120 Hz), high γ (120-250Hz)). Blue dots: binned data points; black line: best linear fit; gray shaded area: 95% C.I. F and p-values are indicated in the main text. Behavioral performance, y-axis: ratio between hit and miss trials in %. Distance between the decoded attentional spotlight (AS) and actual target presentation location, x-axis: normalized distance.

In order to better understand which frequency bands held the most reliable spatial information, the above described correlation analysis is reproduced for each independent functional LFP frequency band (Fig. 3C). Overall, correlations are weak for the lower frequency bands and increase for the higher frequency ranges (Fig. 4C: δ: r^2^= 0.18, F= 0.0, p-value < 0.005/ θ: r^2^= 0.13, F= 0.0, p-value < 0.005/ α: r^2^= 0.26, F= 0.0, p-value < 0.005 / low β: r^2^= 0.13, F= 0.0, p-value < 0.005/ high β: r^2^= 0.17, F= 0.0, p-value < 0.005/ low γ: r^2^= 0.29, F= 0.0, p-value < 0.005 / mid γ: r^2^ = 0.42, F= 0.0, p-value < 0.005 / high γ: r^2^ = 0.40, F= 59.3, p-value < 0.005).

**Figure 4:**
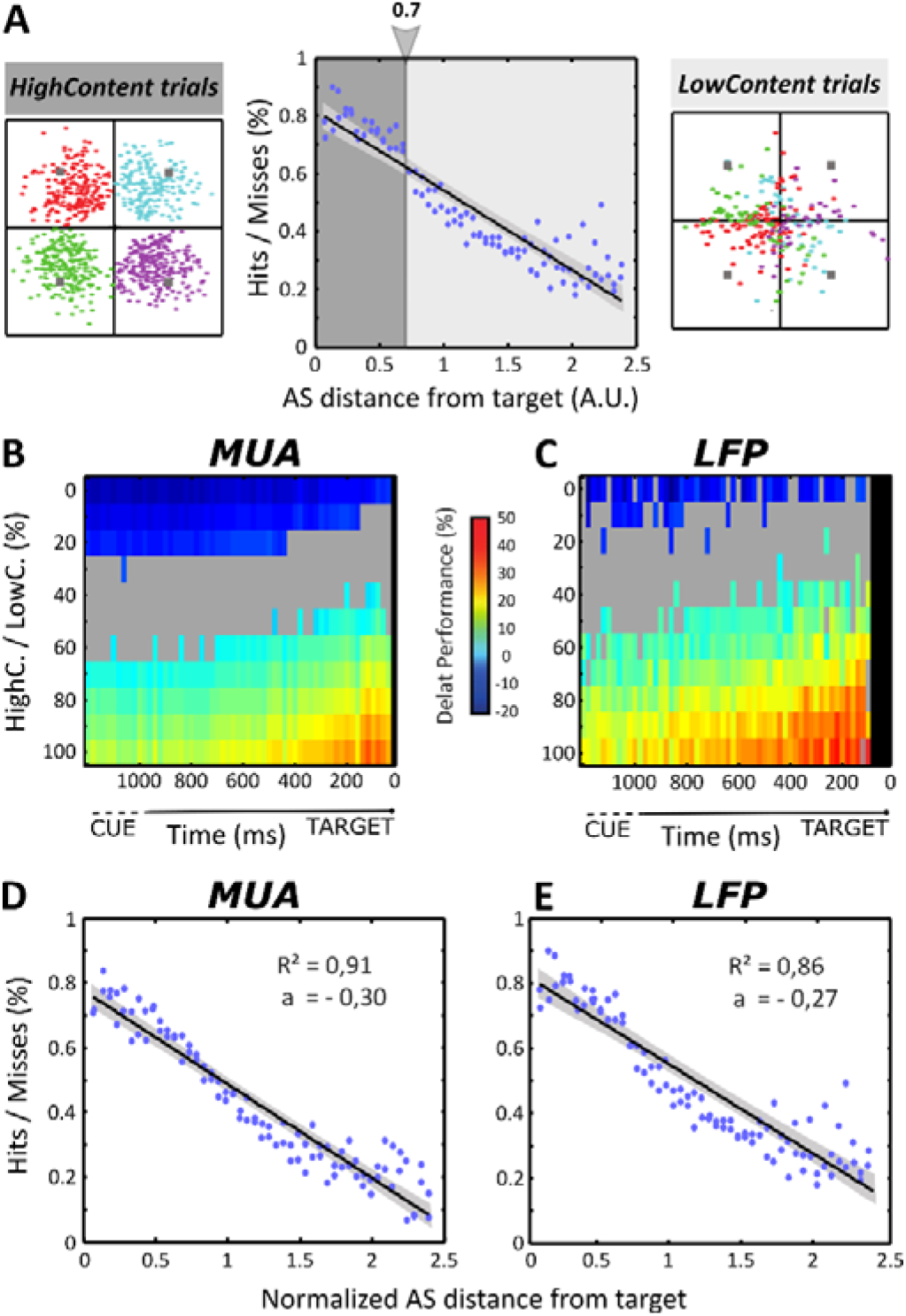
Two-step decoding procedure improves correlation between behavioral performance & distance between the attentional spotlight and the target location. (A) Following the first decoding step, hit trials can be subdivided into HighContent and LowContent trials based on how close decoded attentional spotlight is to the actual target location (mid panel). HighContent trials consistently fall in the cued quadrant (left panel) while LowContent trials don’t. (B-C) Following the second decoding step, the higher the proportion of HighContent trials in the training set, the higher the attention decoding accuracy on novel trials. This improvement in attention decoding accuracy is more marked when decoding from LFP signals (C) than from MUA signals (B) (HighContent trials (HighC.); LowContent trials (LowC.); Shaded gray are: no significant difference in performance as assessed by Wilcoxon rank sum test; Shaded black are: time intervals excluded due to absence of HighContent trials for 5 sessions. (D-E) This two-step decoding procedure improves the correlation between overt performance and the distance of the decoded attentional spotlight (AS) to the target location (Higher R^2^, steeper slope) for both MUA signals (D) and LFP signals (E).

These analyses bring about two important observations. First, spatial attention LFP-based decoding correlates with behavior to the same extent as MUA-based decoding. Second, this is mostly due to the gamma frequency LFP power content.

### Optimizing (x,y) access to attentional spotlight using a two-step decoding procedure

From the above correlation between decoded attentional spotlight distance to expected target location and hit rate, we observe that for a proportion of hit trials, the decoded (x,y) attentional spotlight is estimated close to the expected target location, while for the rest of the trials, the decoded attentional spotlight is estimated far away from the expected target (Fig. 4A). Based on the observation that decoded location accounts for behavior, we reasoned that when training our decoder on hit trials, we are actually training it on suboptimal conditions, presenting it with both trials in which attention is close to the expected target location, and trials in which attention is farther away. We thus here define two different categories of trials: HighContent trials (Fig. 4A), defined by decoded attentional spotlight to expected target distance inferior to 7° and LowContent trials (Fig. 4A), defined by decoded attentional spotlight to expected target distance superior to 7°. Running the decoder on varying proportions of HighContents trials relative to LowContent trials critically impact spatial attention decoding performance. Using an optimal 100% HighContent trials training set from MUA signal leads to an average increase in decoding of 27% (s.e.= 0.8%) between 10 ms to 600 ms pre-target averaging window sizes and an average increase of 18.7% (s.e.=0.3%) between 600 ms to 1200 ms pre-target (Fig. 4B, Wilcoxon rank sum test, p-value < 0.05). Using an optimal 100% HighContent trials training set from LFP signal leads to an average increase in decoding of 34% (s.e.=0.9%) and 25% (s.e.=0.7%), respectively for the short and long pre-target averaging window sizes (Fig. 4B, Wilcoxon rank sum test, p-value < 0.05). This effect was particularly striking for smaller window sizes. A significant increase of performances with respect to a randomly distributed dataset is observed for a minimum threshold of 70% of HighContent trials in the MUA training set and 50% in the LFP training set (Fig. 4B, Wilcoxon rank sum test, p-value < 0.05). In addition, and in contrast with what is described in figure 2, decoding accuracy increment is most marked for shorter than for longer time intervals. Overall, the higher the HighContent trials rate, the higher the gain is in attention classification performance. This indicates that prior selection of a spatial information rich training dataset is crucial to optimize access to prefrontal attentional encoding and further improves classification performances on remaining trials.

In contrast, the higher the LowContent trials rate, the higher the loss in overall spatial attention decoding performance. A training set of 100% LowContent trials leads to a drastic reduction of decoding performance compared to a randomly distributed training set both for MUA signals (Fig. 4B, −13% (s.e.=0.4%) between 10 ms to 600 ms and −16% (s.e.=0.1%) between 600 ms to 1200 ms, Wilcoxon rank sum test, p-value < 0.05) and LFP signals (Fig. 4B, −10% (s.e.=0.8%) and −12% (s.e.=0.6%), Wilcoxon rank sum test, p-value < 0.05). A significant decrease in spatial attention decoding accuracy as compared to a random training dataset is observed for MUA (resp. LFP) training sets starting from 80% or more LowContent trials (resp. 90%, Fig. 4B, Wilcoxon rank sum test, p-value < 0.05). Thus, LowContent trials are detrimental to spatial attention decoding accuracy.

Importantly, the positive effect of HighContent trials on decoding performance is more marked for LFP signals than MUA signals (Wilcoxon rank sum test, p-value <0.005). Moreover, LFP signals are less impacted by the lower ratios of HighContent trials over LowContent trials than MUA signals - thus resulting in a lower decrease in decoding performance (Wilcoxon rank sum test, p-value <0.005). In other words, while the two-step decoding improves attention decoding accuracies, this impact is more pronounced on LFP signals than on MUAs.

Functional validity of this two-step decoding procedure implies that exclusive training on HighContent trials, whether in MUA or LFP signals, maximizes the correlation between the decoded attentional spotlight to expected target distance and behavioral performance (Astrand et al., 2016). We thus trained a decoder using only HighContent trials and tested it on misses and remaining HighContent trials (50% of hit testing trials) and LowContent trials (50% of hit testing trials) to simulate a balanced proportion of hit trials categories and misses. As expected, HighContent trials based decoding increases the linear relationship between attentional spotlight to target distance and behavioral performance. Specifically, in the MUAs, r^2^ value increased from 0.48 (Fig. 3A linear regression: r^2^ = 0.48, F= 85.8912, p-value <0.05) to 0.91 (Fig. 4C, linear regression: r^2^ = 0.91, F= 962, p-value <0.005), and correlation slope becomes markedly more steep (Fig. 4C, linear regression: a=-0.3, vs. Fig. 3A linear regression: a=-0.12). In the LFPs, r^2^ values increase from 0.65 (Fig. 3B linear regression: r^2^= 0.65, F= 174, p-value <0.005) to 0.86 (Fig. 4D linear regression: r^2^= 0.86, F= 569, p-value <0.005), and correlation slope also becomes steeper (Fig. 4D linear regression: a=0.27, vs. Fig. 3B linear regression: a=0.10). Overall, this thus confirms the functional validity of this two-step decoding procedure, both for MUA-based decoding of spatial attention, as well as for LFP-based decoding of spatial attention. Crucially, we demonstrate that using spatial information enriched trials (i.e. HighContent trials) allows to better account for the relationship between observed behavior and the (x,y) decoded attentional spotlight.

## Discussion

In this manuscript we report for the first time high decoding accuracy of the (x,y) location of the attentional spotlight based on prefrontal LFP signals, to a comparable degree to that achieved from MUA signals. We show that both decoded information (MUA and LFP signals) are predictive of behavioral content and that LFP attention-related information is maximal in the gamma band. In addition, we show that selecting maximally attention-informative trials (HighContent trials) during the decoding procedure strongly improves the correlation between our MUA and LFP based decoding and behavioral performance, thus further refining the functional relevance of this decoding of the (x,y) locus of the attentional spotlight. This improvement is more marked for LFP signals than for MUA signals, suggesting that LFP signals may contain other sources of task-related variability than spatial attention information. In the following, these findings are discussed in the light of the current literature.

### Decoding attentional information from LFP signals

The neural bases of spatial attention in the prefrontal cortex have been extensively studied based both on neuronal spiking activity, local field potentials and interferential studies (Ibos et al., 2013; Buschman and Miller, 2007; Wardak et al. 2006). In recent years, this accumulated knowledge has set the grounds for real time decoding of attention both from invasively recorded spiking activity (Astrand et al., 2014, 2016, 2020; Farbod Kia et al., 2011; Gaillard et al., 2020; Tremblay et al., 2015b) and non-invasive brain signals (Andersson et al., 2012, 2011; Thiery et al., 2016; Van Gerven and Jensen, 2009). Attentional decoding methods from MUA signal have made substantial progress, moving from the classification of attention into subspace sectors to the actual decoding of the (x,y) position of the attentional spotlight (Astrand et al., 2016; Tremblay et al., 2015). However, progress has been much slower in the decoding of attention from non-invasive MRI or EEG signals. Decoding of attention from LFP signals and developing novel decoding strategies on this type of signals can be considered as an intermediate step towards improving the decoding of attention from less invasive signals.

To our knowledge, only one study to date has addressed the decoding of spatial attention from prefrontal LFP, based on a four spatial quadrant classification approach (Tremblay et al., 2015a). Here, we report, for the first time the real-time tracking of the (x,y) attentional spotlight locus from prefrontal LFPs. Crucially, we show that the extracted (x,y) locus of the attentional spotlight is highly predictive of the behavioral performance, such that the closer the attentional spotlight to the target presentation location, the higher the correct detection rate. In contrast, the further away the attentional spotlight to the target presentation location, the higher the miss rate. This is important in two ways. First, this result validates the behavioral relevance of the decoding procedure, describing a direct behavioral relationship between where the decoded attentional spotlight is in space relative to where the target is presented and the detection rate of the subject. Second, this indicates that very much like has been described from MUA-based attentional spotlight tracking, the LFP-based attentional spotlight is highly dynamic and explores space even when cued towards a specific location. Indeed, the LFP-based decoded attentional spotlight is not anchored at the expected target location following cue presentation, but can be more or less close to this task-relevant location, in spite of the fact that behavioral performance is enhanced when the attentional spotlight is closest to the cued location.

As previously described Tremblay et al., (2015a), we confirm that attention-related information is maximal in the LFP gamma frequency band (above 30Hz, and maximally between 60 and 120Hz). Attention-related information can still be extracted above chance in lower LFP frequency bands, though at much lower accuracies. These results are in agreement with the description of the contribution of gamma frequency bands to attentional processes (Chalk et al., 2010). From a methodological point of view, there is no benefit in classifying attention-related information from gamma frequency bands. Indeed, full spectrum LFP decoding accuracy is higher compared to LFP gamma frequency band decoding accuracies. This result suggests that attention related information in the multiple frequency bands is not fully redundant.

The correlation between decoding and behavior is further enhanced using the two-step decoding procedure that we introduce here and that is discussed below. This latter point is crucial for neurofeedback and cognitive brain-machine interfaces (Andersen et al., 2010; Astrand et al., 2014; Enriquez-Geppert et al., 2017; Jiang et al., 2017; Ordikhani-Seyedlar et al., 2016), where one wants to work with information of maximal behavioral relevance. Interestingly, Salari et al., (2014) demonstrate a modulation of perception by a neurofeedback manipulation based on EEG gamma power. This is possibly in agreement with our observation that gamma frequency contains high attention-related information. However, these studies are based on direct modulation of surface gamma power, independently from behavioral performance or a global extraction of attentional spotlight locus. Our approach allows to track the dynamic attentional spotlight with a high temporal resolution (down to 30ms). We expect this type of approach to provide subjects with more informative and reliable neurofeedback to work on.

### Exploiting attention dynamics to improve real-time attention decoding accuracies

The fact that the attentional spotlight is extremely dynamic (Gaillard et al., 2020) suggest that not all hit trials are equivalent. Indeed, we observe that some hit trials take place when the attentional spotlight is successfully located where the target appears and other hit trials in contrast happen when attention is far away from target presentation location. This has a direct impact on decoding performances. The more space sampled, less stable the information in the neuronal population, thus impairing resulting decoding performance. On the contrary, a trial with less exploration and a more stable spotlight will lead to a stable neuronal information and more accurate decoding. Based on these observations, we reasoned that training our classifier on all of these hit trials is suboptimal as compared to training the classifier on hit trials in which attention was properly oriented. We thus use a first decoding step to identify such good trials (i.e., high attention-related information content or HighContent trials) and specifically use them to train the decoder on a second decoding round. This significantly increases the attention decoding accuracies. Several points need to be noted. First, as expected from our initial hypothesis, the higher the proportion of HighContent trials used for the training the higher the relative gain in decoding accuracies. Strikingly, for both MUA and LFP signals, decoding improvement is higher when considering short time interval compared to longer time window. This observation could be explained by the fact that the longer the time window, the more attention is expected to explore the target position, this both on HighContent and LowContent trials. Quite importantly, this increment in decoding accuracies was more marked for the LFP decoding than for the MUA decoding. This possibly indicates that LFP signals multiplex attention related information with other sources of information, contributing to LFP signal variability, and that are more prevalent on LowContent than on HighContent trials. Last but not least, this two-step decoding procedure drastically improves the correlation between the (x,y) attentional spotlight real-time estimate and behavioral performance, whether from MUA or LFP signals. In other words, the decoded attentional spotlight better explains behavior, both as assessed from the strength of the correlation and from its slope.

Overall, our work presents two major advances in the field of real-time access to the attentional spotlight locus. First we demonstrate that this spotlight location can be estimated from both MUA and LFP signals. Second, we introduce a novel two-step decoding method that further enhances the behavioral relevance of the decoded attentional spotlight. Most crucially, our work illustrates the tremendous benefit of adapting machine learning strategies to the specific functional properties of the cognitive function under study.

## Acknowledgments

S.B.H was supported by ERC Brain3.0 #681978, ANR-11-BSV4-0011 & ANR-14-ASTR-0011-01, LABEX CORTEX funding (ANR-11-LABX-0042) from the Université de Lyon, within the program Investissements d’Avenir (ANR-11-IDEX-0007) operated by the French National Research Agency (ANR). C.D.S.F. and C.G. were supported by ERC Brain3.0 #681978. We thank research engineer Serge Pinède for technical support and Jean-Luc Charieau and Fidji Francioly for animal care. All procedures were approved by the local animal care committee (C2EA42-13-02-0401-01) and the Ministry of research, in compliance with the European Community Council, Directive 2010/63/UE on Animal Care.

## Contributions

Conceptualization, S.B.H. C.D.F. and C.G.; Data Acquisition, S.B.H.H., F.D.B.; Methodology, C.D.F. C.G and S.B.H.; Investigation, C.D.F. C.G and S.B.H; Writing – Original Draft C.D.F. C.G and S.B.H; Writing – Review & Editing, C.D.F. C.G and S.B.H; Funding Acquisition, S.B.H.; Supervision, S.B.H.

## Code availability

The code that supports the findings of this study is available from the corresponding author upon reasonable request. The code is still being used for other purposes and cannot be made publically available at this time.

